# Potential geographic distribution of the Tiger Mosquito *Aedes albopictus* (Skuse, 1894) (Diptera: Culicidae) in current and future conditions for Colombia

**DOI:** 10.1101/2020.03.13.990440

**Authors:** Emmanuel Echeverry-Cárdenas, Carolina López-Castañeda, Juan D. Carvajal-Castro, Oscar Alexander Aguirre-Obando

## Abstract

In Colombia, little is known on the distribution of the Asian mosquito *Aedes albopictus*, main vector of dengue, chikungunya, and Zika in Asia and Oceania. Therefore, this work set out to estimate its current and future potential geographic distribution under the Representative Concentration Paths (RCP) 2.6 and 8.5 emission scenarios by 2050 and 2070, using ecological niche models. For this, predictions were made in MaxEnt, employing occurrences of *A. albopictus* from their native area and South America and bioclimatic variables of these places. It was found that, since its invasion to Colombia, *A. albopictus* is present in 47% of the country, in peri-urban (20%), rural (23%), and urban (57%) areas between 0 and 1800 m, with Antioquia and Valle del Cauca being the departments with the most registries. The current estimation suggests that *A. albopictus* is distributed in 96% of the territory up to 3000 m (p < 0.001). Additionally, by 2050 and 2070, below RCP 2.6, its distribution could diminish to nearly 90% including altitudes of 3100 m, while below RCP 8.5 it would be < 60% increasing its distribution up to 3200 m. These results suggest that, currently in Colombia, *A. albopictus* is found throughout the country and climate change could diminish eventually its area of distribution, but increase its altitudinal range. In Colombia, surveillance and vector control programs must focus their attention on this vector to avoid complications in the national public health setting.

## INTRODUCTION

The tiger mosquito, *Aedes albopictus* (Skuse, 1894) (Diptera: Culicidae), presents vector competence for at least 26 arboviruses and some filiarial nematode worms (1, 2). In continents, like Asia and Oceania, *A. albopictus* is the main vector for dengue, chikungunya, and Zika (3–6). In America, it is not considered as the prime vector for these arboviruses, however, sporadically, it has been found infected naturally with dengue in countries, like the United States (North America), Colombia, and Brazil (South America) (7–9). Additionally, the tiger mosquito could present an ecological niche similarity with *Aedes aegypti* (10), the primary vector for dengue, chikungunya, and Zika in this continent, and whose presence in Colombia encompasses 90% of the territory up to 2.300 m. Currently, for these three arboviruses, no efficient vaccines exist yet (11–13).

The tiger mosquito is native to tropical, subtropical, and temperate forests of Asia and the islands of the western Pacific (14). In these zones, favorable conditions for its development for the aquatic immature phases are estimated at water temperature between 26 and 32 °C, while the adults require environmental temperature ranging between 25 and 31 °C and relative humidity > 70%. In addition, it has been detected in temperatures >40 °C and below 17 °C its survival is notably affected (15–17). In unfavorable environmental conditions, this species presents the diapause phenomenon (diminished metabolism to very low rates of energy expenditure and subsequent inactivity) in the development of its eggs, which has permitted its dispersal at latitudes with temperate and seasonal climates, beyond its range of native distribution (18–20). This invasion has been largely facilitated by human activities, like passive transport via maritime, land, or air cargo (21). Due to the aforementioned, it has been suggested that *A. albopictus* exposes high ecological plasticity, considered among the 100 most invasive species in the world (21, 22).

Chronologically, regarding its global invasion, *A. albopictus* was first registered outside its native distribution range in Europe, specifically in Albania in 1979 (23). Thereafter, the first populations of this species were registered in America; initially, in the center, in Trinidad and Tobago in 1983 (24), then in the north, in the United States in 1985 (25), and in the south, in Brazil in 1986 (26). In this last part of the continent, particularly in Colombia, the tiger mosquito was first registered in Leticia (Amazon, on the border with Tabatinga, Brazil) in 1998, in a suburban area with abundant vegetation (27). Since then, it has been registered in 52 locations of 12 departments of the 32 that make up the country (28). However, vast zones still remain where its presence is unknown and given its vector competence, it is necessary to recognize them to include them in the surveillance and control programs in public health.

One way of complementing the lack of knowledge of the distribution of *A. albopictus* in Colombia is through ecological niche modeling (ENM). This tool enables characterizing the fundamental niche of a species and then estimate its potential geographic distribution from registries of presence and environmental variables (29–32). Given the relevance of the ENM for public health, these have been used previously to estimate the potential distribution of mosquitoes of medical importance belonging to the *Haemagogus* (33), *Culex* (34), *Anopheles* (35) and *Aedes* (10, 36) genera. Particularly for *A. albopictus*, its potential distribution has been estimated in Australia (37), western Europe (38), the United States (39), Mexico (40), Guatemala (41), and globally (10,42,43).

Furthermore, climate change could influence directly on the geographic distribution of invasive mosquitoes. Taking into consideration the different gas emission scenarios (*i.e*., RCP 2.6 or RCP 8.5), investigations conducted until now suggest that the geographic distribution of *A. albopictus* could vary significantly in the long term, which would imply that the viral diseases transmitted by this vector could disperse to new places in the country (10,14,21,44,45). Due to the aforementioned, it is necessary to better understand the current distribution of *A*. *albopictus* and its likely future variations in Colombia. Therefore, this work sought to estimate and quantify the current potential geographic distribution of this vector in Colombia and identify the effect of climate change on its distribution under RCP 2.6 and 8.5 emission scenarios by 2050 and 2070 by using the ENM approach.

## MATERIALS and METHODS

### Study area

The Republic of Colombia is located in northeastern South America and borders geographically with the republics of Venezuela, Brazil, Peru, Ecuador, and Panama. Additionally, it has coastal zones on the Caribbean and on the Pacific Ocean. Its continental extension is of 1,141,748 Km^2^ and its political-administrative division comprises 32 departments (46).

### ENM and estimation of M

In eastern Asia, the native distribution range for *A. albopictus* is concentrated in the biomass: tropical and subtropical rain forest, tropical and subtropical dry forest, temperate forest, and mixed forest. Starting from the aforementioned, registries of occurrence of the tiger mosquito in said biomass were used to characterize its accessible area (M); see ahead for more details. Then, spatial and temporal transfers were made toward South America to identify its potential distribution areas. From each transfer, estimations corresponding to the area of Colombia were extracted to describe the current and future potential distribution of the tiger mosquito.

### Data of *A. albopictus* presence

From a published literature review, reports available in the Colombian National Health Institute (47) and the Global Biodiversity Information Facility (GBIF) database (48) two sets of occurrence data were formed. The first, compiled the occurrences of the native range of the tiger mosquito available in the GBIF and those collected by Kamal *et al*., (10). The second set gathered the occurrences of *A. albopictus* in South America. Of these, for occurrences in Colombia, information was extracted, like altitude, type of coverage, and area of location. Thus, the first and second datasets were conformed initially by 2,085 and 3,414 registries, respectively. Data cleaning was conducted, excluding registries without spatial geo-referencing, with geo-spatial problems, duplicate presence, and multiple presence in a single pixel, at a resolution of 2.5 min (∼5 Km^2^) (10,39,42). For this, the *raster* 3.0-7 (49), *rgdal* 1.4-8 (50), *dismo* 1.1-4 (51) and *usdm* 1.1-18 (52) libraries of R (53) were used. After the data filtering, the first and second datasets were consolidated with 1,328 and 3,406 occurrences, respectively.

### Climate data

From the WorldClim database v. 2.0, 21 environmental variables were downloaded with a spatial resolution of 2.5 min (10,39,42), whose values are based on averaged data since 1970 to 2000 (54). These variables were submitted to two analyses to define their inclusion in the calibration of the models. First, the contribution of each variable was determined through the Jackknife test generated in MaxEnt, maintaining those whose accumulated contribution added to 95%. Then, with the variables selected a Spearman R correlation was conducted. Variables highly correlated positively (R > 0.8) or negatively (R < – 0.8) were discarded for the ENM.

To assess the potential distribution of the species within a context of climate change, the variables resulting from prior analyses were downloaded from the Climate Change, Agriculture and Food Security - CCAFS (55) platform, with values estimated by the HadGEM2-ES model for 2050 and 2070, for RCP 2.6 and 8.5 emission scenarios. The HadGEM2-ES model, developed by the Hadley Center (UK), is one of the most adequate to analyze future projections in tropical areas of South America (56–60). Within this model, different climate scenarios are projected formulated by the Intergovernmental Panel on Climate Change (IPCC), known as Representative Concentration Paths (RCP), which estimate distinct greenhouse gas emission levels and CO_2_ over time. Among them, there is RCP 2.6 based on a gas emissions peak (∼ 421 ppm), being the scenario with lowest effects on climate, and RCP 8.5 based on continuous increase of gas emissions (∼ 936 ppm), considered the scenario with the most drastic climate effects (61). All the layers of the variables selected were adjusted to the extension of M defined and from South America using QGIS v.3.4.0 (62), for its later use in the estimations described ahead.

### Geographic distribution estimations

Three contexts were proposed to analyze the potential geographic distribution of *A. albopictus* in Colombia, and in each its latitudinal and altitudinal variation were identified. For the first context, the first dataset was used, together with the layers of the environmental variables under the current conditions cut to the native and South American extension. In the two remaining contexts, the potential effects of climate change were estimated on the distribution of the tiger mosquito in Colombia by 2050 and 2070, through the emission paths RCP 2.6 and 8.5 for each period, respectively.

All the estimations were made through the maximum entropy algorithm, implemented in the MaxEnt software v.3.4.1 k (63). This algorithm was used due to its high accuracy when estimating distribution areas, allowing to calibrate the models through datasets of different sizes, determining the contribution of each environmental variable in the estimations performed; it may be used for predictions in multiple spatial and temporal scales and only requires presence data to conduct the estimations (64–66). For each scenario proposed, 10 replicas were executed per 1000 iterations, using a logistic output format. For future estimations, the parameters “*Do Clamping*” and “*Extrapolation*” were deactivated to avoid extrapolations in the extreme values of the ecological variables (non-analog climates) (10).

The estimations obtained in MaxEnt were reclassified in a binary format to distinguish potential distribution areas of the tiger mosquito. For this, the threshold approach corresponding to the lowest environmental suitability value was followed associated with known presence registries, considering an emission (E) value of 0.2 (67–70). Finally, the potential distribution area was quantified in all the scenarios.

### Validation of the model

This work only evaluated the performance of the model under current conditions, given that the behavior of *A. albopictus* is unknown upon eventual future climate scenarios. To do so, the metric of the area under the curve (AUC) was considered as estimated in MaxEnt. Additionally, to obtain greater support on the performance of the model, the AUC significance level was determined through a partial analysis of the Receptor’s Operational Characteristics (partial ROC), employing the second dataset (69, 71). Analyses of partial ROC were carried out on the Niche Toolbox platform (72), where the E parameter was adjusted to 0.2 per 1000 iterations. As criterion to evaluate the model’s significance, it was considered that AUC values with p > 0.05 indicate that the estimations made are not better than those generated by a random model, while AUC with p < 0.05 indicates that the predictions estimated are better than those obtained from a random model (36, 69).

## RESULTS

Since the first registry, in 1998 in Colombia, *A. albopictus* has been registered in 52 locations of 15 departments, between 0 and 1800 m. Information was gathered for 45 locations, of which 27 had information about the location of the capture sites. The seven locations not collected correspond to poorly detailed INS information or to personal communications to other authors. The departments with more occurrences registered were Antioquia (24.5%) and Valle del Cauca (22.5%). In addition to this, the urban area is where the presence of the tiger mosquito has prevailed (57%), followed by rural areas (23%) and peri-urban areas (20%) (Figure 1). In urban areas, the tiger mosquito has been associated principally with relicts of forests immersed in the urban matrix (Table 1).

**Figure 1.**
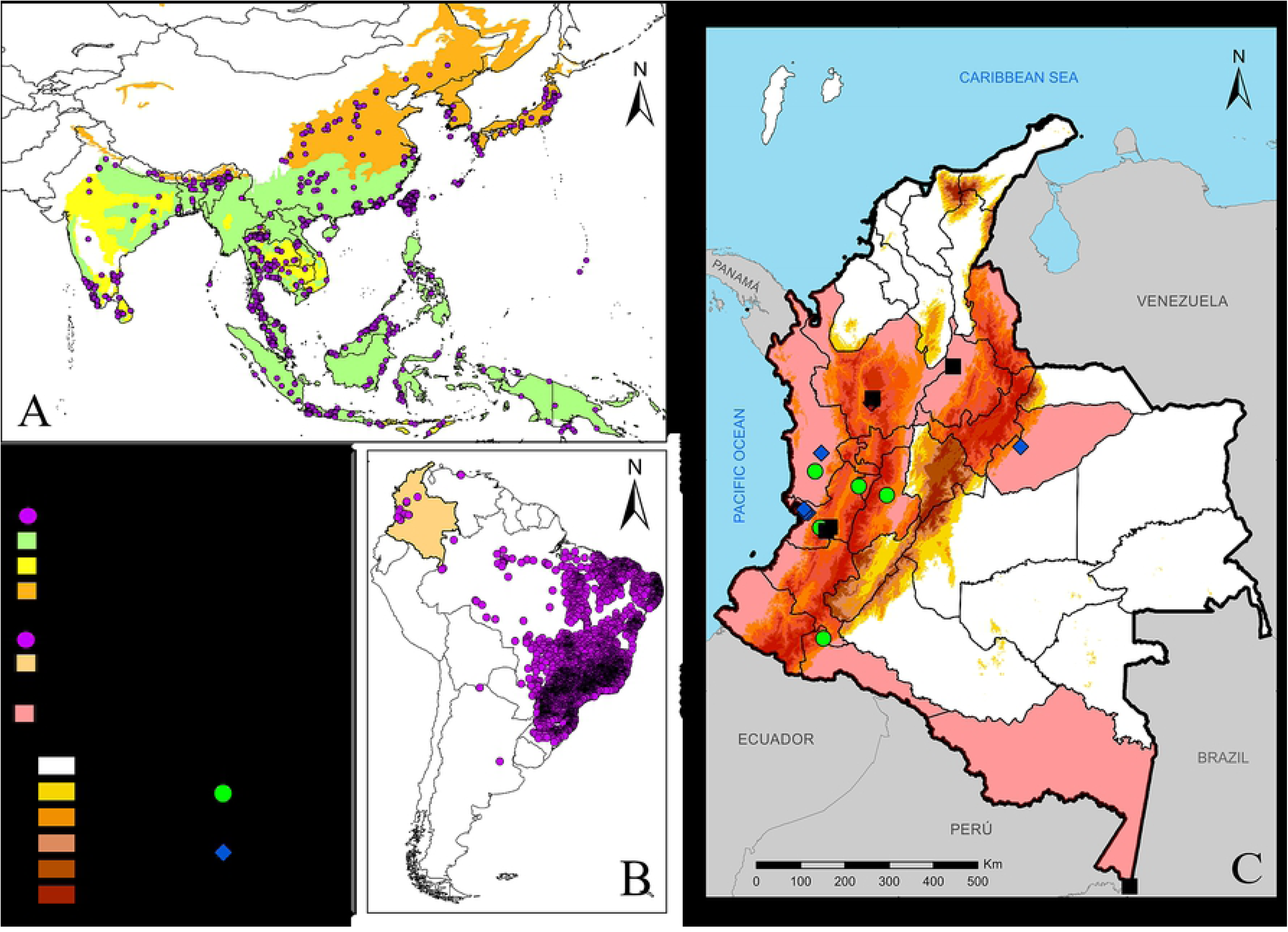
Occurrences of *A. albopictus* in: **A.** Native area (first dataset), **B.** South America (second dataset), and **C.** Colombia, employed in the ENM.

**Table 1.**
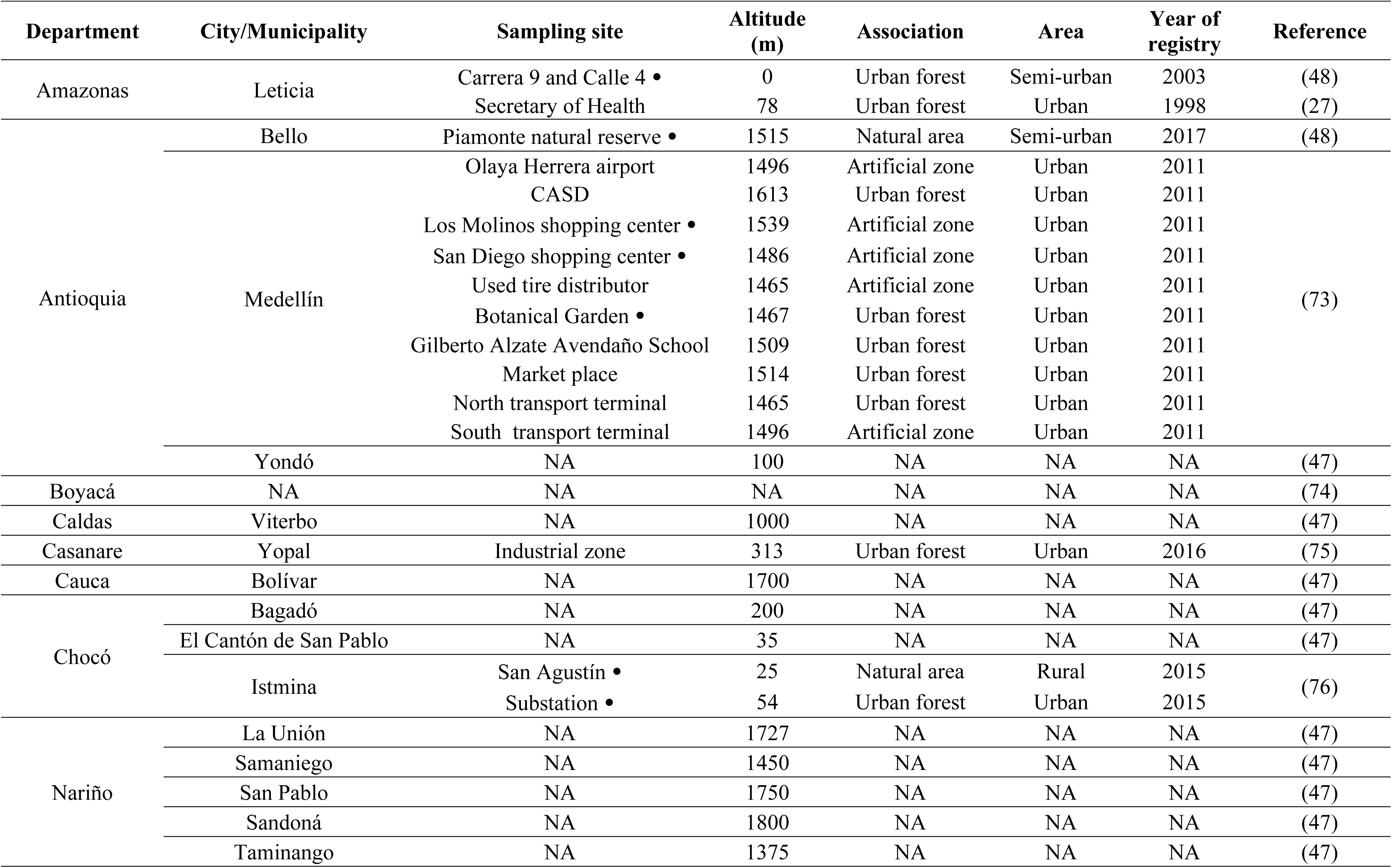
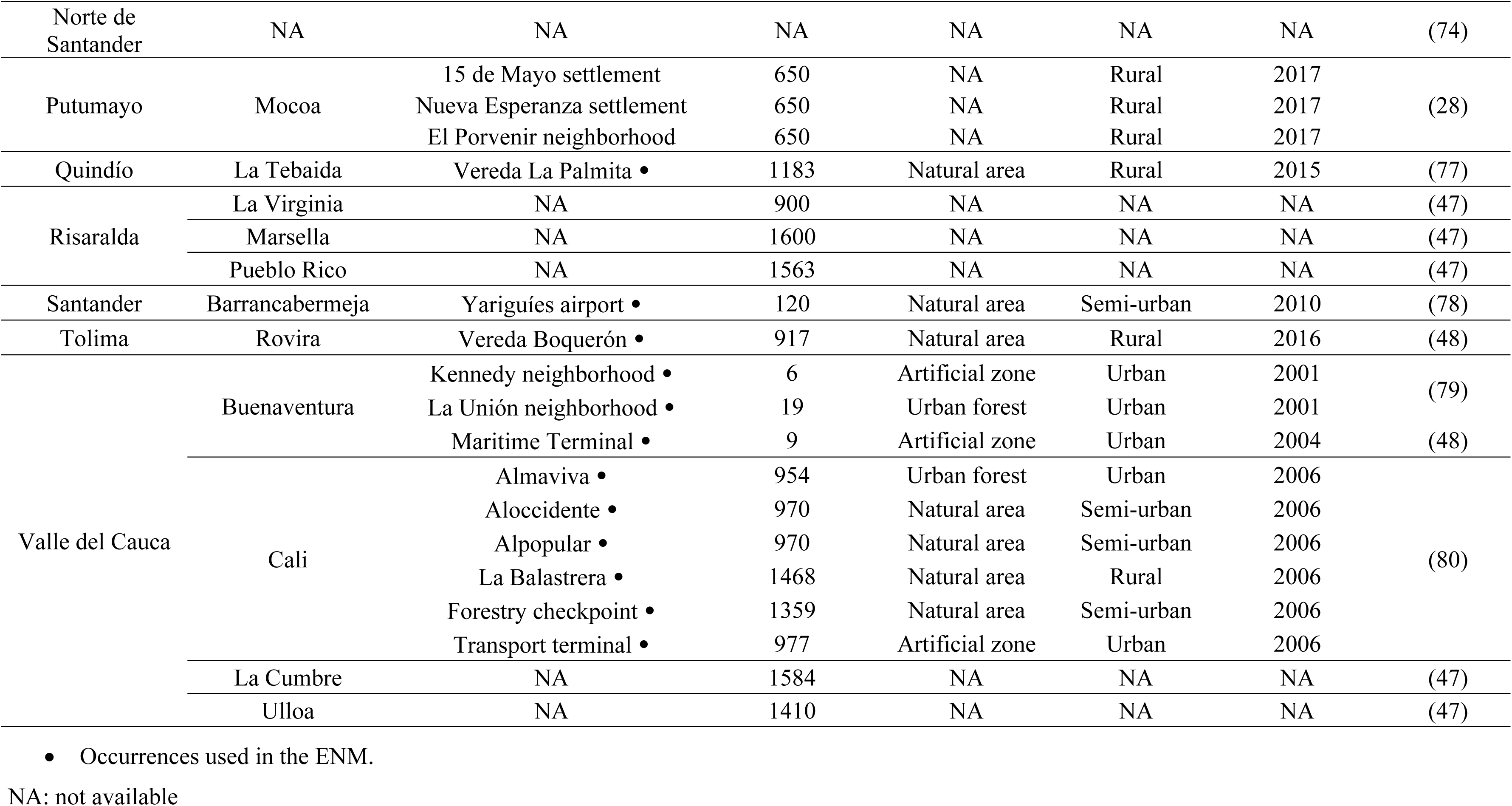
Registries of *Aedes albopictus* in Colombia and its border limits.

Table 2 presents the environmental variables used to calibrate the ENM, including variables of mean annual temperature and annual precipitation; although a high correlation was present, due to their importance in the life cycle of *A. albopictus*.

**Table 2.** Climate variables used in the ENM for the tiger mosquito in Colombia.

| Variables | Unit |
| --- | --- |
| Mean annual temperature | °C |
| Range of daily temperatures | °C |
| Isothermality | % |
| Annual precipitation | mm |
| Precipitation of the rainiest month | mm |
| Precipitation of the driest month | mm |
| Precipitation of the rainiest quarter | mm |
| Precipitation of the warmest quarter | mm |

For Colombia, results of predictions of *A. albopictus* currently estimated its presence in 96.14% of the territory (Table 3) in all the departments, including altitudes up to 3.000 m (Figure 2). The AUC metric estimated in MaxEnt was 0.9, while the partial ROC supported statistically the predictions (p < 0.001).

**Figure 2.**
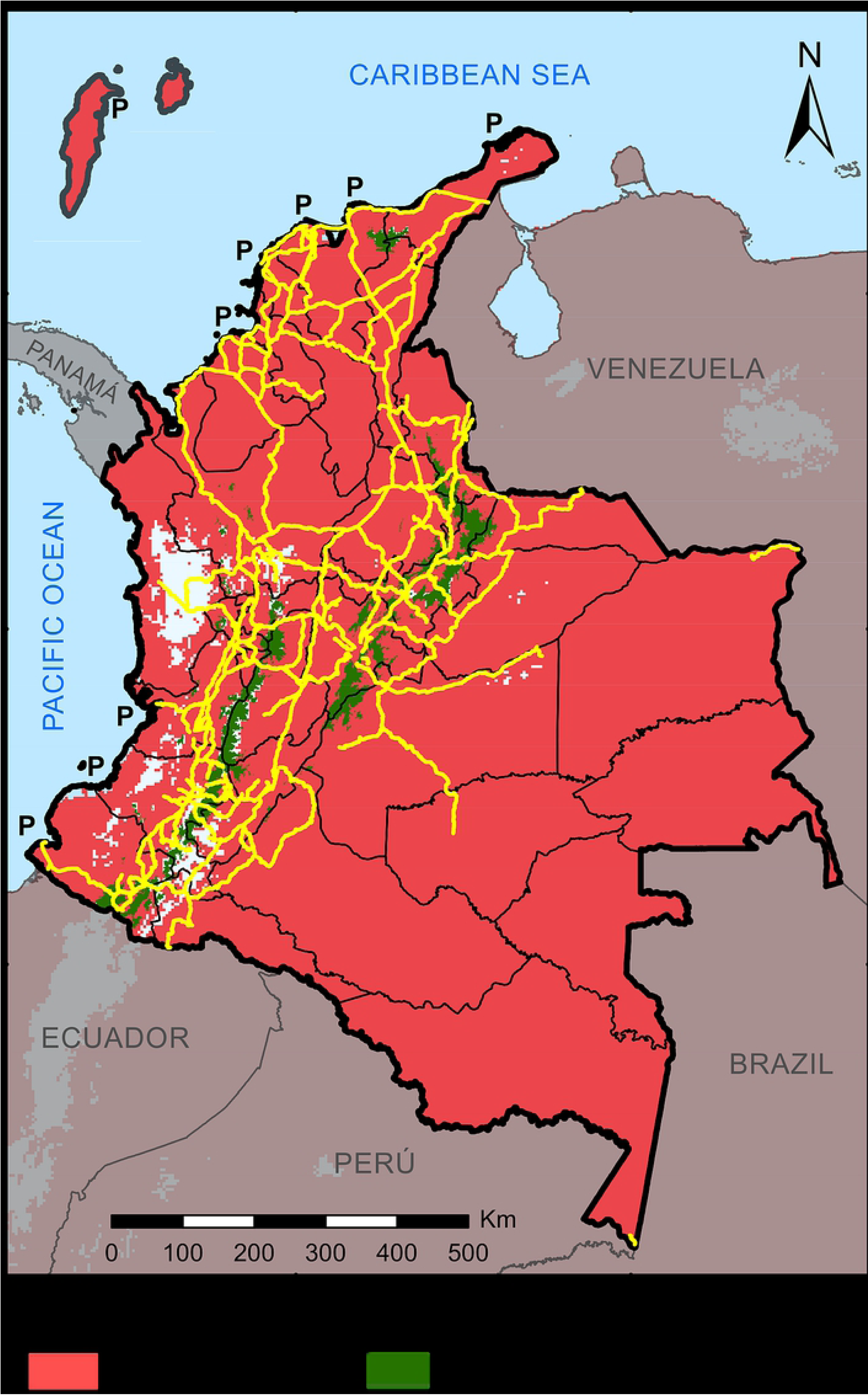
Potential geographic distribution of *A. albopictus* under current conditions in Colombia.

**Table 3.**
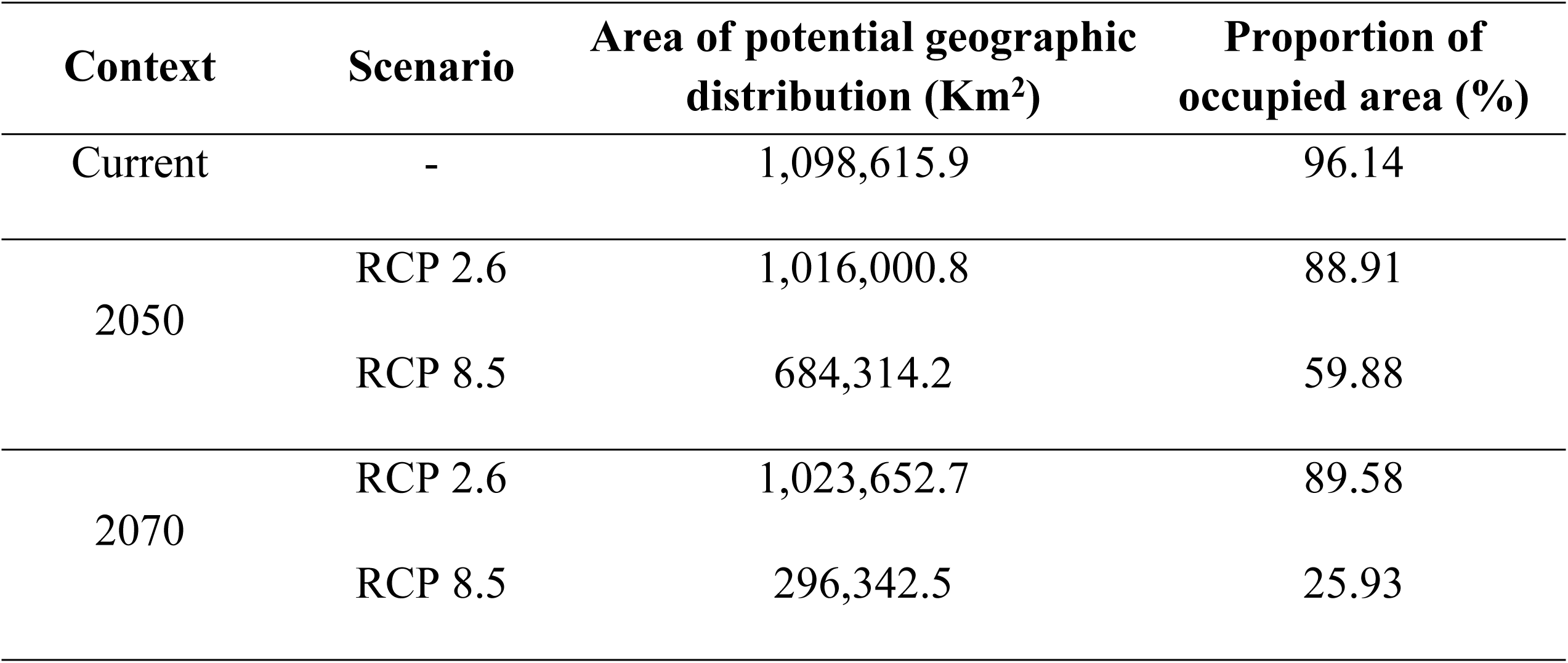
Areas of current and future potential distribution of *A. albopictus* in Colombia.

Predictions of *A. albopictus* within a context of climate change for 2050 and 2070 estimated that the departments of Nariño, Cauca, Huila, Quindío, Risaralda, Caldas, Cundinamarca, and Boyacá could have the same distribution observed currently. Under the RPC 2.6 emission scenario, the tiger mosquito had the same distribution pattern in which it could continue present in over 85% of the country (Table 3) and increase its distribution range up to 3100 m for both years. On their behalf, in departments, such as Chocó, Valle del Cauca, Cauca, Vichada, Santander, Cesar, Bolívar, La Guajira, and San Andrés y Providencia greater decrease could occur in the potential area with respect to current values (Figure 3). Additionally, under the environmental conditions of the RCP 8.5 emission scenario by 2050 and 2070, *A. albopictus* could eventually broaden its altitudinal range up to 3200 m. By 2050, environmental conditions could provoke a decrease of its distribution (Table 3) in the departments of La Guajira, Magdalena, Atlántico, Bolívar, Sucre, Córdoba, Cesar, western Santander, eastern Norte de Santander, Eastern Tolima, Chocó, western Valle del Cauca, western Cauca, Arauca, Casanare, Vichada, Meta, Guainía, and Guaviare. Besides these departments, by 2070, the area of potential distribution could also diminish in peripheral zones of Antioquia, Vaupés, Caquetá, Putumayo, and Amazonas, where its distribution would be restricted to the departments associated with the Andes mountain rage principally, like Nariño, central-eastern Cauca, central-eastern Valle del Cauca, Huila, western Tolima, Quindío, Risaralda, Caldas, central-southern Antioquia, Cundinamarca, Boyacá, eastern Santander, and central-western Norte de Santander, besides buffer zones of the Sierra Nevada of Santa Marta, to the north of the country (Figure 4).

**Figure 3.**
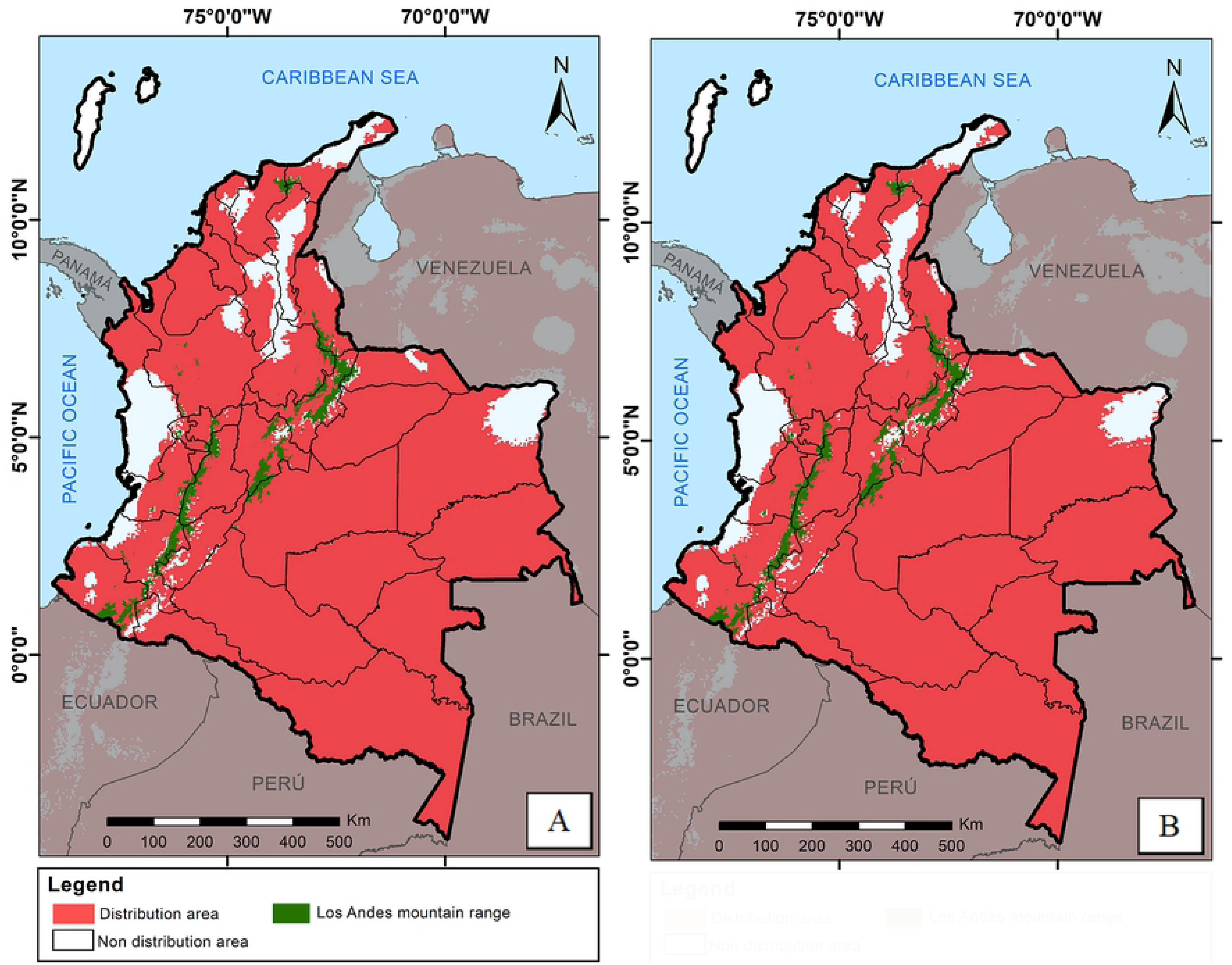
Potential geographic distribution of *A. albopictus* within a context of climate change for: **A.** 2050 and **B.** 2070, under the RCP 2.6 emission scenario.

**Figure 4.**
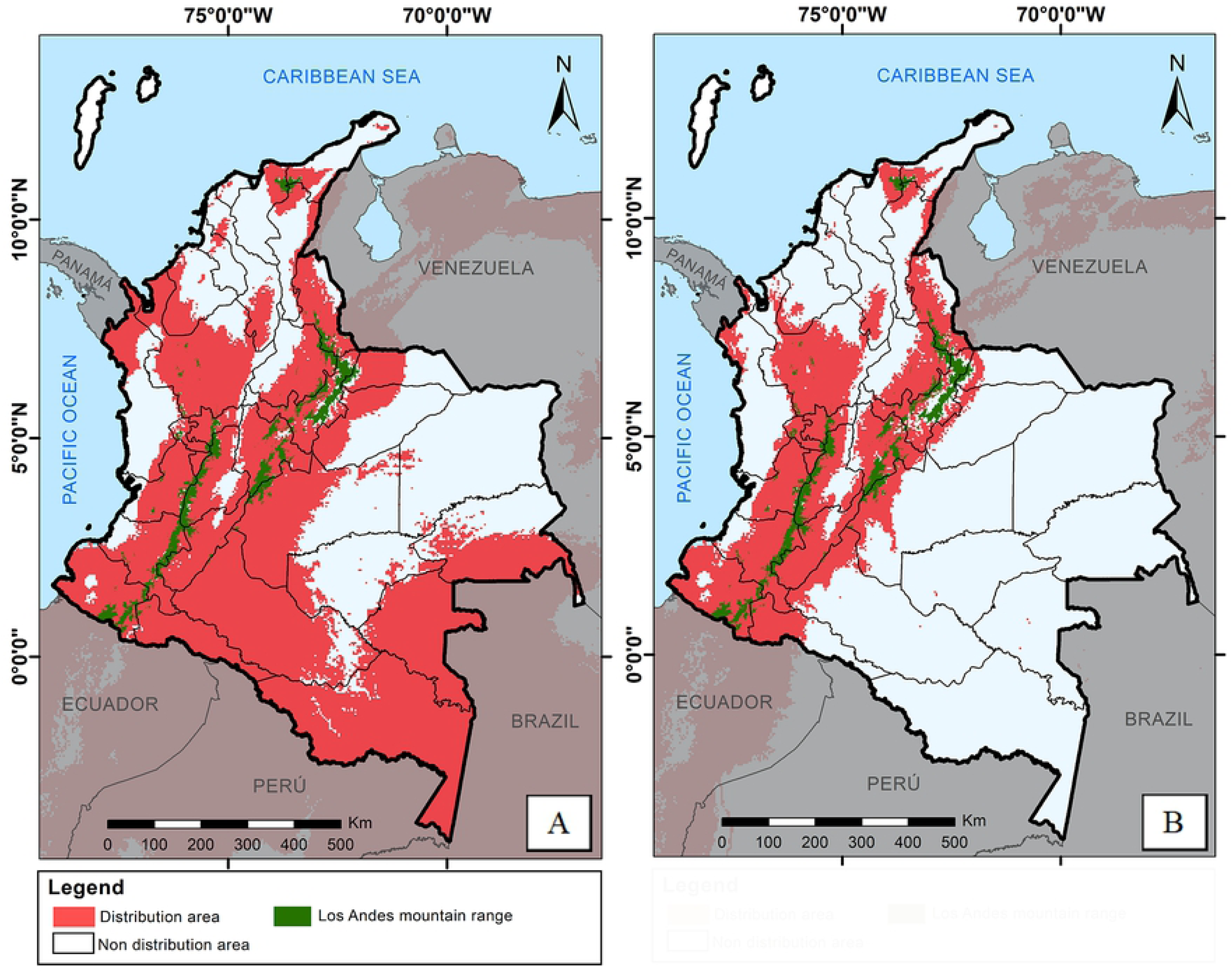
Potential geographic distribution of *A. albopictus* within a context of climate change for: **A.** 2050 and **B.** 2070, under the RCP 8.5 emission scenario.

## DISCUSSION

Current estimates suggest that *A. albopictus* could have a broad distribution in Colombia. By 2050 and 2070, under the RCP 2.6, its distribution could diminish up to nearly 90% including altitudes of 3100 m, while under the RCP 8.5 it would be close to 60%, increasing its distribution up to 3200 m. It has been observed that the invasion of this mosquito to other countries started in the coastal zones (81) and thereafter disseminated to their interior (82, 83). In this sense, we can hypothesize that ports to the Pacific Ocean of Buenaventura (Valle del Cauca), Guapi (Cauca) and Tumaco (Nariño) (84), through which 50% of commercial imports enter the country in ships, most of them from Asia (native place to *A. albopictus*), could have played an important role in its initial invasion to the country. Furthermore, it should not be discarded that maritime ports located on the Atlantic Ocean (Caribbean) in the departments of Sucre, Bolívar, Atlántico, Magdalena, La Guajira, and San Andrés y Providencia (85), where official reports of this vector are still not available, also could have facilitated its invasion. Nevertheless, in both cases, genetic evidence is required to support these hypotheses. Added to this, climate conditions of all the coastal departments mentioned (86) are similar to the conditions registered in its native area, thereby, favoring its survival and reproduction (87). An increase has been observed in coastal zones of cases of diseases transmitted by vectors, principally of *Anopheles* and *Aedes* genera, due to El Niño and La Niña climate phenomena, which have favored increments of artificial oviposition sites (water tanks, containers, etc.) or natural sites (plants, puddles, etc.) and, consequently, increasing the population size of the vectors and the probability of arbovirus transmission (88–92).

Upon establishing the populations of the tiger mosquito in the departments with coastal zones, land passive transport may have also played an important role in its distribution to the rest of Colombia, as noted in other parts of the world (93). High roadway connectivity, as well as the national vehicular flow between the center, west and north of the country, and international connections with western Venezuela – where registries already exist of *A. albopictus* (94), would permit rapid invasion of the tiger mosquito to new departments (81– 83,95,96).

In Santander, Antioquia, Quindío, Caldas, Risaralda, and Tolima, where the tiger mosquito has been registered in 19 locations (47,48,73,77,78), a current broad distribution was also estimated. The vast geographic and environmental heterogeneity (mix of natural and urban areas) and urban-rural transitions of these departments, similar to those of its native area, increases the availability and diversity of microhabitats, as well as the number of breeding sites in which the tiger mosquito could develop its immature stages and increase quickly its population size (73). In addition to this, the country’s human population and the 492 mammal species reported (97, 98) represent potential food sources and, thereby, subsistence for the tiger mosquito (10, 14). Furthermore, in these places *A. aegypti* is widely disseminated up to 2300 m, together with the circulation of dengue, chikungunya, and Zika (99) for which this species is the principal vector in America. Due to this, the role of *A. albopictus* in the transmission of these arboviruses cannot go unnoticed given the panorama mentioned and this vector should be included in surveillance and control strategies of said diseases, given that new alternatives to control *A. aegypti* are being implemented in this continent. Among said strategies, we can highlight the use of transgenic mosquitoes (known as Release of Insect Carrying a Dominant Lethal Gene *--* RIDL; mosquitoes released seeking to eliminate the vector in a particular location) and infected with *Wolbachia pipientis*–WMel lineage (mosquitoes with refraction to arboviruses transmitted by *A. aegypti*) (100, 101). Which is why, if in any zone of the country with presence of both species we could suppress or establish populations of *A. aegypti* refractory for dengue, chikungunya, and Zika, the tiger mosquito could assume the function of principal vector of these arboviruses (102–104).

In 17 departments of Colombia the presence of *A. albopictus* has not been reported; however, predictions indicate that it would also be present in such, probably because such areas comply with the environmental requirements for its distribution (11). Therefore, it becomes necessary to guide studies to detect this vector in these locations and apply adequate strategies to prevent its dissemination.

In addition, increased temperature, sea level, and precipitation variability are some effects brought by climate change, Therefore, some places in which now *A. albopictus* could be present, in the future would not have adequate conditions for its permanence (105). Nonetheless, in mountainous zones of Colombia where temperatures are currently cold and act as an ecological restriction for invading arthropods (106), variations in temperature could favor the establishment of the tiger mosquito even in altitudes above those that have been currently registered (up to 1800 m) by 2050 (up to 3100 m) and 2070 (up to 3200 m) (18,19,47).

Under the RCP 2.6 emission scenario, a decrease could occur in the distribution area of the tiger mosquito by 2050 and 2070 in some departments characterized historically by high temperatures, like Vichada and Guajira (20) and those mostly affected by El Niño phenomenon, like coastal zones (Chocó, Valle del Cauca, and Cauca).

Under the RCP 8.5 scenario, we suggest that the environmental conditions could change drastically by 2050 and 2070, which would provoke a considerable decrease in the distribution of the tiger mosquito in the country with respect to current values, as hypothesized globally (43). In this order of ideas, this vector’s distribution could be limited in most of the departments associated with the Andes mountain range, which would maintain favorable conditions for its survival.

## CONCLUSION

Currently, *A. albopictus* is distributed in 96% of Colombia, including altitudes up to 3,000 m, being the country’s environmental conditions, the food sources, and passive transport possible key factors for its invasion to new departments where it still has not been registered. Moreover, the effects of climate change by 2050 and 2070 could generate increase in its altitudinal range up to 3200 m and affect the presence of the tiger mosquito in the country’s coastal, plains, and jungle zones, but could remain principally in the Andean departments. Finally, greater attention should be paid to this potential vector in Colombia, given that it has a similar niche as that of *A. aegypti*, as well as vector competence for dengue, chikungunya and Zika, which would complicate public health in the country.

## ACKNOWLEDGMENTS

The authors thank Universidad del Quindío for the financial support of Project 885. Likewise, gratitude is expressed to Doctors Jonny E. Duque-Luna and Andrés Arias-Alzate for their valuable contributions to this manuscript.

## CONTRIBUTIONS BY THE AUTHORS

Design and construction of the models: EEC, CLC, JDCC and OAAO.

Execution of the models: EEC.

Interpretation of the results: EEC, CLC and OAAO.

Manuscript drafting: EEC and OAAO. All the authors read and approved the final version of the manuscript.

